# Spatial top-down proteomics for the functional characterization of human kidney

**DOI:** 10.1101/2024.02.13.580062

**Authors:** Kevin J. Zemaitis, James M. Fulcher, Rashmi Kumar, David J. Degnan, Logan A. Lewis, Yen-Chen Liao, Marija Veličković, Sarah M. Williams, Ronald J. Moore, Lisa M. Bramer, Dušan Veličković, Ying Zhu, Mowei Zhou, Ljiljana Paša-Tolić

## Abstract

**Background:** The Human Proteome Project has credibly detected nearly 93% of the roughly 20,000 proteins which are predicted by the human genome. However, the proteome is enigmatic, where alterations in amino acid sequences from polymorphisms and alternative splicing, errors in translation, and post-translational modifications result in a proteome depth estimated at several million unique proteoforms. Recently mass spectrometry has been demonstrated in several landmark efforts mapping the human proteoform landscape in bulk analyses. Herein, we developed an integrated workflow for characterizing proteoforms from human tissue in a spatially resolved manner by coupling laser capture microdissection, nanoliter-scale sample preparation, and mass spectrometry imaging.

**Results:** Using healthy human kidney sections as the case study, we focused our analyses on the major functional tissue units including glomeruli, tubules, and medullary rays. After laser capture microdissection, these isolated functional tissue units were processed with microPOTS (microdroplet processing in one-pot for trace samples) for sensitive top-down proteomics measurement. This provided a quantitative database of 616 proteoforms that was further leveraged as a library for mass spectrometry imaging with near-cellular spatial resolution over the entire section. Notably, several mitochondrial proteoforms were found to be differentially abundant between glomeruli and convoluted tubules, and further spatial contextualization was provided by mass spectrometry imaging confirming unique differences identified by microPOTS, and further expanding the field-of-view for unique distributions such as enhanced abundance of a truncated form (1-74) of ubiquitin within cortical regions.

**Conclusions:** We developed an integrated workflow to directly identify proteoforms and reveal their spatial distributions. Where of the 20 differentially abundant proteoforms identified as discriminate between tubules and glomeruli by microPOTS, the vast majority of tubular proteoforms were of mitochondrial origin (8 of 10) where discriminate proteoforms in glomeruli were primarily hemoglobin subunits (9 of 10). These trends were also identified within ion images demonstrating spatially resolved characterization of proteoforms that has the potential to reshape discovery-based proteomics because the proteoforms are the ultimate effector of cellular functions. Applications of this technology have the potential to unravel etiology and pathophysiology of disease states, informing on biologically active proteoforms, which remodel the proteomic landscape in chronic and acute disorders.

## Background

Investigation of proteoforms,^1^ which includes post-translational modifications (PTMs), splice-isoforms, and amino acid variants on a canonical protein sequence has gained increased attention in recent years. Proteoforms hold great potential to become more precise molecular biomarkers than their respective proteins, transcripts, and genes.^2, 3^ Mass spectrometry (MS) is a powerful tool to directly characterize proteins; however, routine bottom-up proteomics (BUP) cannot fully capture the molecular identity of proteoforms.^4^ Due to proteolytic digestion within BUP, proteoform level information is commonly lost or is not trivial to reconstruct.^5^ Top-down proteomics (TDP) overcomes this by avoiding enzymatic digestion and allowing for the analysis of intact proteoforms. An exemplary use of TDP was the recent completion of the Blood Proteoform Atlas, where roughly 30,000 proteoforms were detected across 1690 human genes. Many of these proteoforms demonstrated high cell-type specificity.^6^ Furthermore, PTMs are known to regulate cellular functions. For example, phosphorylation plays important roles in apoptosis and cell growth,^7^ and conformation and activity of proteins are widely controlled by glycosylation.^8^ The identity and roles of all proteoforms have yet to be fully characterized, therefore there is a present need to further contextualize which proteoforms effect function.^3^ As such, advanced TDP methods must be developed and applied for confident and comprehensive proteoform characterization.

Historically, the sensitivity of TDP has been problematic,^9^ especially for higher molecular mass proteins.^10^ Confident identification of proteoforms therefore has required large sample inputs for high resolution accurate mass measurements and efficient fragmentation. This has traditionally limited TDP to bulk-scale tissue analyses and large quantities of cultured cells. However, recent advances in MS instrumentation and bioinformatic tools have significantly improved TDP sensitivity and coverage.^11^ This progress has enabled the field of TDP to move towards the detection and discrimination of proteoforms directly from tissues in a spatially resolved manner. Several direct analysis techniques have been deployed at near cellular resolution for the analysis of proteins from tissue. Some notable advancements use mass spectrometry imaging (MSI) such as matrix assisted laser desorption ionization (MALDI),^12, 13^ nanospray desorption electrospray ionization (nanoDESI),^14–16^ and liquid extraction surface analysis (LESA).^17, 18^ While many of these techniques are routinely used for probing metabolites, lipids, and peptides within functional tissue units (FTUs),^19^ and in single cell analyses,^20, 21^ the spatial analysis of intact proteoforms remains challenging in both proteoform coverage and spatial resolution.

Notably, translation of these direct analysis techniques into preclinical studies is particularly challenging,^22^ where the lack of routine chromatographic or gas-phase separation with various matrix effects lowers the sensitivity and dynamic range in comparison to traditional liquid chromatography tandem mass spectrometry (LC-MS/MS). In the case of MALDI-MSI analyses, low charge states of the generated ions lead to inefficient fragmentation which poses difficulty in proteoform identification.^23^ To address the challenge in detecting and quantifying proteoforms, the field has made considerable progress over the past decade.^24^ Spatial proteomics has also been broadly expanded using methods such as microLESA,^25^ liquid microjunction microextraction (LMJ),^26^ parafilm-assisted microdissection (PAM),^26^ and laser capture microdissection (LCM)^27^ to circumvent some challenges of MSI. Nanodroplet processing in one-pot for trace samples (nanoPOTS), and its companion microPOTS, have enabled routine single cell proteomics and BUP processing of dissected tissues.^28–30^

In a previous study, we demonstrated the coupling of nanoPOTS sample preparation with TDP enabling the identification of over 150 proteoforms along with various PTMs from roughly 70 cultured HeLa cells.^31^ This technology was further extended to spatial TDP by integrating nanoPOTS with LCM to identify an average of 509 proteoforms per 100,000 µm^2^ dissection in brain tissue.^10^ Here we further demonstrate LCM-TDP of representative regions-of-interest, which benefits from the chromatographic separation and enables deeper coverage of proteoforms. In tandem with MALDI-MSI we utilize the accurate mass database of proteoform from LCM-TDP to map proteoforms with high spatial resolution and throughput (or field-of-view). Herein, we highlight an advanced workflow for comprehensive spatial proteomics, where LCM-microPOTS-TDP and MALDI-MSI are integrated for high-confidence identification, near-cellular localization, and label-free quantitation of proteoforms within human kidney tissue.

## Methodology

### Experimental Design and Motivations

The generalized workflow for processing fresh-frozen tissues for tandem LCM-microPOTS-TDP and MALDI-MSI is illustrated within **Figure 1**. A critical consideration for confident proteoform identification from MALDI-MSI is the use of serial or near serial sections from the same tissue block for TDP, where bulk analyses from different donors can result in less accurate databases for MALDI-MSI annotations. In depth protocols for each step reported below have been deposited in the protocols.io repository.^32, 33^ The workflow generally benefits from efficient capture of dissected regions of interest within microdroplets on manufactured microPOTS chips **(Figure 1a)**. This enables minimal protein losses from iterative transfers and sample processing prior to LC-MS/MS analyses and allows quantitative TDP analyses of singular FTUs. However, the throughput for LC-MS based TDP is low, where roughly a dozen dissected regions can be analyzed within a day. Here MALDI-MSI presents utility, with routine processing of serial sections only requiring a few hours. Importantly, a custom ultrahigh mass range (UHMR) Orbitrap **(Figure 1e)** enables the detection of proteoforms at 1 to 2 Hz, allowing for broad profiling of entire tissue sections within 6 to 24 hours (field-of-view dependent) with a near cellular or cellular scale probe (≤ 175 um^2^). This permits the profiling of several dozen FTUs. Proteoforms within these FTUs can be relatively quantified leveraging a strong overlap between the several hundred proteoforms confidently identified by LC-MS/MS and the several dozen proteoforms detected by MALDI-MSI. Herein, healthy human tissues sections were used to exemplify the potential of preclinical applications, which often requires an in-depth analysis of precious samples.

**Figure 1:**
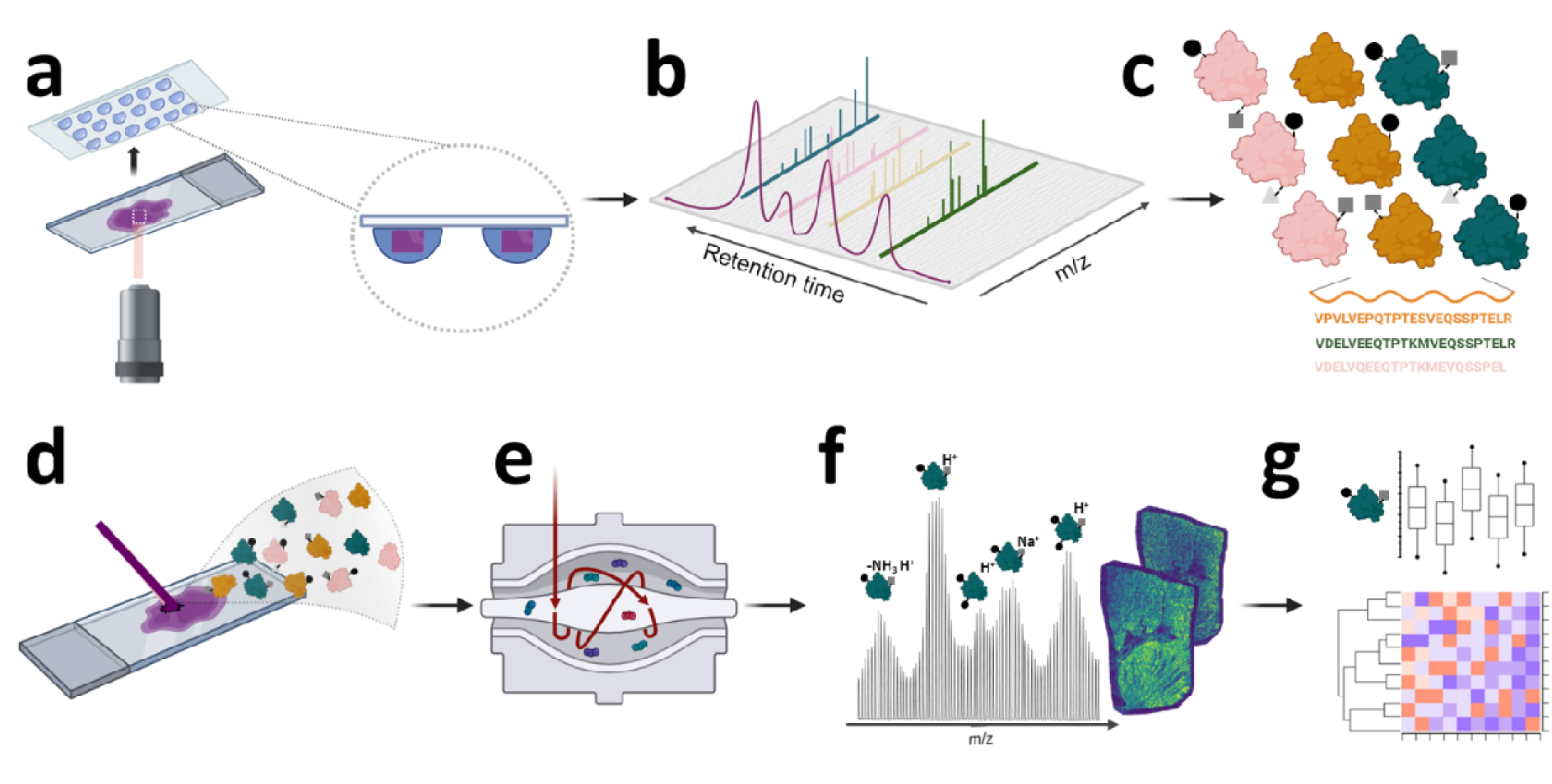
Overall workflow for LCM-microPOTS-TDP and MALDI-MSI. **(a)** LCM process where regions of interest are dissected from the tissue mounted on polyethylene naphthalate (PEN) membrane slides and are catapulted into microdroplets on microPOTS chips for TDP; **(b)** The isolated samples in microdroplets undergo protein extraction followed by LC-MS/MS to **(c)** generate quantitative proteoform databases. **(d)** Serial sections on ITO slides are prepared for MALDI-MSI with **(e)** UHMR HF Orbitrap detection enabling high mass resolving power and high mass accuracy which **(f)** is needed to identify intact proteoforms based on isotopic distributions using top-down databases generated from LCM-microPOTS-TDP in **(a-c)**. **(g)** Bioinformatic tools are developed to confidently annotate MALDI-MSI spectra and statistics, clustering, and segmentation are then applied. Portions of this figure were created with BioRender.com.

### Experimental Samples

Tissue sections were obtained from Vanderbilt University Biomolecular Multimodal Imaging Center (BIOMIC) which is a Tissue Mapping Center affiliated with the Human BioMolecular Atlas Program (HuBMAP).^34, 35^ Briefly, the tissues were cryo-sectioned at 10 µm thickness on indium-tin-oxide (ITO) and polyethylene naphthalate (PEN) membrane slides, and tissues were vacuum sealed shipped with desiccant, on dry ice, and stored at −80 °C until analyses. Further metadata regarding the donor can be found on the HuBMAP data portal as described in the data availability section.

### Laser Capture Microdissection and Sample Preparation

A full step-by-step protocol for the processing of tissues via microPOTS is deposited in the protocols.io repository.^33^ Briefly, an injection molding company, ProtoLabs produced the polypropylene (PP) microPOTS chips used for these experiments. The tissue samples were fixed in 70 % ethanol for 1 min, followed by sequential dehydration in 95 % ethanol and 100 % ethanol for 1 min each. Regions of interest were isolated from the kidney sections mounted on PEN membrane slides using a PALM MicroBeam laser microdissection system (Carl Zeiss MicroImaging, Munich, Germany). Tissue voxels with an area of 100,000 µm^2^ were excised from 10 µm thick tissue sections for six replicates of three different FTUs including glomeruli, tubules, and medullary rays. Each microPOTS well had a diameter of 2.2 mm and was preloaded with 2 µL of DMSO as the capture liquid. DMSO was evaporated by heating the chip at 70 °C and protein extraction was performed by adding 2 µL lysis buffer in each well, containing 2.5 units/µL benzonase nuclease, 2 mM MgCl_2_, 10 mM TCEP, 0.2 % DDM and 4 M urea in 50 mM ABC. The mixture was incubated for 1 hr at 37 °C. Following incubation, the sample was acidified by adding 500 nL of 5 % formic acid into each well and dried thoroughly in a vacuum chamber where the chips were stored at −20 °C until LC-MS/MS analysis.

### Liquid Chromatography Mass Spectrometry Analysis

A step-by-step protocol for TDP LC-MS/MS is deposited in the protocols.io repository.^33^ Briefly, custom trap columns (150 µm i.d., 4 cm long) and analytical columns (100 µm, i.d., 50 cm long) were packed with C2 particles (SMTC2MEB2-3-300, Separation Methods Technologies, Newark, DE) in-house. LC separations were performed using a dual pump NanoAcquity system (186016032, Waters, Millford, MA) equipped with mobile phase A (MPA; 0.2 % FA in water) and mobile phase B (MPB; 0.2 % FA in ACN). For desalting, the samples were loaded over a period of 10 min, and were washed with 95 % MPA at a flow rate of 5 µL/min. The analytical gradient was started at 90% MPA and linearly ramped to 40% MPA over 90 min, where the gradient was then ramped to 90 % MPA over 10 min at a flowrate of 3 µL/min.

An Orbitrap Fusion Lumos Tribid mass spectrometer (Thermo Scientific, San Jose, CA) was operated in intact protein and full profile mode with data-dependent acquisition. For MS1 acquisitions, the instrument was set with the following parameters: a default charge state of 10, resolution of 120K at *m/z* 200, maximum injection time of 500 ms, 4 microscans, mass range from *m/z* 500 to 2000, a 250% normalized AGC target of 1E6, and 15V of in-source fragmentation. For MS2 acquisitions, the instrument was set to dynamically exclude precursor ions 1 time within 30 s period, precursors were isolated with a 2 Da window for CID with fixed collision energy of 35 %, 10 ms activation time, and 0.25 activation Q. A resolution of 120K at *m/z* 200, maximum injection time of 500 ms, mass range from *m/z* 400 to 2000, 2 microscans, and 4 data dependent scans was used. All data were uploaded to the HuBMAP Data Portal for visualization and inspection.

### Liquid Chromatography Mass Spectrometry Data Analysis

A step-by-step protocol for processing TDP LC-MS/MS datasets is deposited in the protocols.io repository.^33^ Briefly, instrument files converted to .mzML were deconvoluted using TopFD.^36^ These exported files were analyzed within the TopPIC Suite (v.1.4.13.1)^37^ and TDPortal (v.4.0.0)^38^. TopPIC spectra were searched against UniProtKB with both SwissProt and TrEMBL sequences (downloaded on June 29, 2019, containing 20,352 reviewed sequences). Default settings were used for TDPortal analysis of the same datasets. TopPIC proteoform spectrum matches (PrSMs) were then imported into the TopPICR companion package for improved identification and quantitation,^39^ where outputs from both TDPortal and TopPIC were combined within the R environment. A quality control filtering step was included to remove samples with very low identification rates. The set threshold was a minimum of 100 proteoform spectrum matches per sample, which led to the exclusion of only two samples (one from medullary dissection and one tubular dissection). All further data analysis steps were performed within the R environment, including median normalization, KNN imputation, and visualization of results.^10, 40^ Imputed data was solely utilized with algorithms that require it, such as PCA analysis and hierarchical clustering. For differential abundance analysis, non-imputed data was used, and statistical testing was performed with Empirical Bayes Method and linear modeling as implemented in the DEP R package (via limma).^41^

### Sample Preparation for Mass Spectrometry Imaging

A step-by-step protocol for sample preparation for intact protein MALDI-MSI is deposited in the protocols.io repository.^32^ Briefly, tissue sections were defrosted and desiccated under vacuum for 30 mins followed immediately by serial washes in fresh solutions of 70% ethanol for 30 s, 100% ethanol for 30 s, Carnoy’s solution (6:3:1 v/v ethanol: chloroform: glacial acetic acid) for 2 mins, 100% ethanol for 30 s, water with 0.2% TFA for 15 s and 100 % ethanol for 30 s in glass Coplin jars, each wash was completed with a volume of 70 mL. Following the washes, the tissue sections were thoroughly dried by a stream of nitrogen gas. An M5 sprayer from HTX Technologies (Chapel Hill, NC) was then used for acidification of the tissue section where a solution of 5% acetic acid (*v/v*) in 50% ethanol was sprayed at a flow rate of 150 µL/min, nozzle temperature of 30 °C, and spray velocity of 1250 mm/min with 10 PSI of nitrogen gas. A “CC” pattern with 3 mm track spacing was used for spraying and a 5 s drying period was applied between each of the five passes. Immediately after tissue acidification and purging of the loop, 15 mg/mL 2,5-dihyroxyacetophenone (DHA) in 90% acetonitrile with 0.2% TFA was deposited using the same M5 sprayer described above. The sprayer was operated at a flow rate of 150 µL/min, nozzle temperature of 30 °C, spraying velocity of 1300 mm/min with 10 PSI of nitrogen gas. The matrix was applied in a “CC” pattern with a 2 mm track spacing. Coverage of DHA was calculated to be ~277 ug/cm^2^. After deposition a recrystallization step was performed with a 5% acetic acid solution in water at 38.5 °C, where the sample was equilibrated for 2 min prior to exposure to vapor for 2 min with a 1 min drying period afterwards.

### Instrumentation for Mass Spectrometry Imaging

The operation of the instrumentation for MALDI-MSI is described in the protocols.io repository.^32^ Briefly, a MALDI source (Spectroglyph, LLC, Kennewick, WA) equipped with an 349 nm Explorer One Nd:YAG laser (SpectraPhysics, Stahnsdorf, Germany)^42^ was mounted on a custom Q Exactive HF Orbitrap MS. The Q Exactive Orbitrap MS was upgraded with Exactive UHMR boards and was operated under custom privileges licenses.^13^ Calibration of the instrument was performed through electrospray of a solution of 2 mg/mL cesium iodide in 50% IPA. MALDI imaging was carried out with a *m/z* range from 3,500 to 20,000 and a lowered noise thresholding setting of 1.5 for acquisitions, resolution of 240k at *m/z* 200, ion accumulation time of 500 ms with a laser frequency of 500 Hz pumped at 1.49 A. This resulted in 250 laser shots per pixel and analysis by light microscopy showed that the laser spot size was roughly 12 µm by 15 µm. The source pressure was regulated to 7.0 Torr with in-source trapping at −300 V.

### Data Processing for Mass Spectrometry Imaging

The data processing and bioinformatics workflow for MALDI-MSI annotations were deposited in the protocols.io repository.^32^ Briefly, FreeStyle (v.1.4, Thermo Scientific, Bremen, DE) was used to view the reduced profile .RAW files. The positional data files (.xml) were exported from the Spectroglyph, LLC source to SCiLS Lab software (v.2021c, Bruker Daltonics, Bremen, DE) for generation of ion images with automatic settings. Processing and root mean square transformation was completed for generation of the ion images. To guide the segmentation of the regions of interest in the SCiLS Lab the tissue sections were stained with periodic acid-Schiff (PAS) after MALDI-MSI experiments with matrix washed off prior. High-resolution bright-filed images were overlayed with the MALDI images for manual segmentation. Mass errors for each proteoform were also reported from a single isotope in the average spectrum, and all SMART annotations are found within captions of all ion images.^43^ Peak-by-Peak (Base Edition, v.2021.8.1.b4, SpectroSwiss, Lausanne, CH) was used to average all pixels and convert spectra to .mzML for further annotation within IsoMatchMS^44^ within a search range of *m/z* 4800 to 16000, a noise threshold of 10 %, a Pearson correlation cutoff of 0.8, and all other parameters left at the default values for intact proteins. No additional adducts (*i.e.*, sodium or potassium) were searched. All samples including the TDP library and pooled replicates were used to generate the MALDI-MSI annotation library.

### Materials and Chemicals

Optima LC-MS grade glacial acetic acid (99.99%), trifluoroacetic acid (TFA, purity 99.5 %), tris(2-carboxyethyl) phosphine (TCEP), n-dodecyl-beta-maltoside (DDM) detergent, protease/phosphatase inhibitor cocktails (catalog 78,430), acetonitrile (ACN), isopropanol (IPA), and water were HPLC grade and were purchased from Fisher Scientific (Fair Lawn, NJ). Ethanol (200 proof) was acquired from Decon Laboratories (King of Prussia, PA), while chloroform, 2’,5’ dihydroxyacetophenone (2,5-DHA), cesium iodide (CsI), magnesium chloride (MgCl_2_), phosphate buffered saline (PBS) and ammonium bicarbonate (ABC) were purchased from Sigma Aldrich (St. Louis, MO). Benzonase nuclease was purchased from EMD Millipore (Darmstadt, Germany).

## Results

### LCM-microPOTS-TDP Profiling of Human Kidney Proteoforms

The routine dissection of FTUs was visualized through a combination of bright-field and fluorescence microscopy as highlighted in **Figure 2**, which permits facile LCM of cortical FTUs in kidney. Additionally, the MALDI-MSI presented was completed prior to LCM of FTUs on a serial section, where we identified distinct proteoform localizations in the medulla directed by ion images. This MSI informed LCM has previously been demonstrated with a metabolome-informed proteome imaging (MIPI) approach.^45^ Here, we applied LCM to dissect six replicates for each of three FTUs from cortical and medullary regions: glomeruli, tubules (both distal and proximal), and medulla (which largely consists of medullary striations, consisting of loops of Henle). A pooled sample representing all anatomical regions was used to create a proteoform database which is referred to as “library” below to aid in annotation of the smaller dissected regions and MALDI-MSI features. In total 616 proteoforms were identified across all samples, with an average of 123 proteoforms annotated per sample. Coefficients of variation (CV) were higher within FTUs relative to bulk bottom-up proteomic analyses, reflecting the challenge of very low input samples with high degrees of heterogeneity (**Supplementary Figure 1**). Binning proteoforms into groups based on data completeness demonstrates at least 80 to 100 proteoform in each FTU have fewer than 50% missing values (**Supplementary Figure 2**). Across all samples, 113 proteoforms had at least 50% complete data and therefore could be utilized in downstream multivariate analyses after imputation.

**Figure 2:**
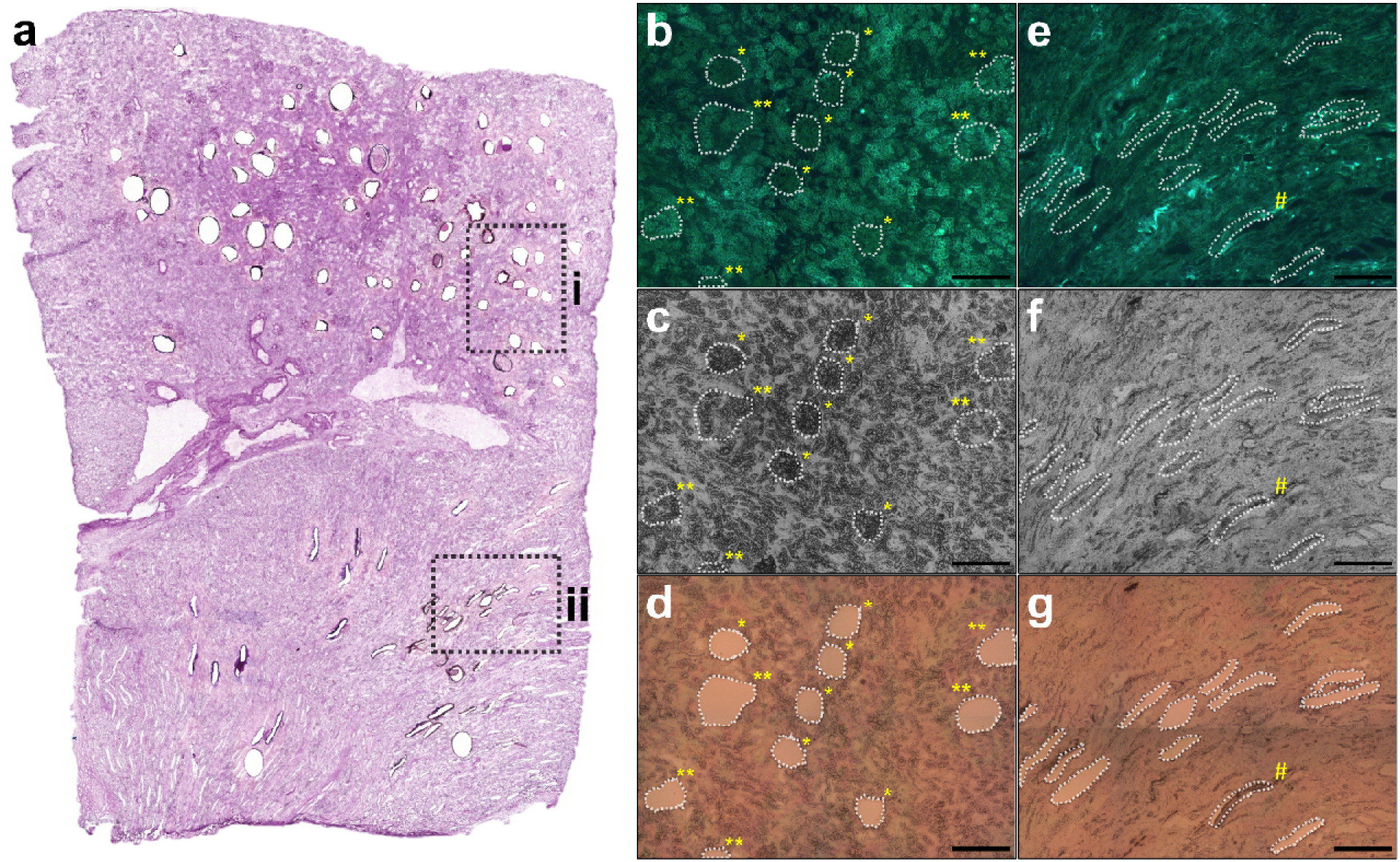
Cortical and medullary dissections from human kidney for LCM-microPOTS-TDP. **(a)** Shows high-resolution image of the histological image taken post LCM processing for library production and dissection of glomeruli, tubules, and medullary rays. The inset region **(i)** represents the region of interest highlighted for **(b)**, **(c)**, and **(d)** where several glomeruli and tubules were dissected and the region is rotated within the zoome visualization, where * denotes dissected glomeruli and ** denotes dissected tubules. The inset region **(ii)** represents the region of interest highlighted for **(e)**, **(f)**, and **(g)** where several areas of the medullary rays were dissected, where # denotes a medullary section which was dissected but not catapulted. **(b)** and **(e)** represent the autofluorescence image before LCM, **(c)** and **(f)** are the grayscale bright-field images before LCM, and **(d)** and **(g)** are the bright-field images after LCM. The scale bar for **(b)**, **(c)**, **(d)**, **(e)**, **(f)**, and **(g)** is 250 µm.

Notably, the pooled library sample, representing the aggregate of all FTUs, appeared to be centered within the first two principal components and equally distant between all different FTUs analyzed **(Figure 3a)**. Differential abundance analysis of medulla compared to tubules showed no significant differences in proteoform abundances, however this is not entirely unexpected as the two FTUs are anatomically and functionally very similar despite being collected from different regions of the kidney (**Figure 3b**). Comparison of glomeruli and medulla showed several mitochondrial proteoforms enriched in medulla (**Figure 3c**), similar to tubules (*vide infra*). Looking at just glomeruli and tubules via PCA shows these two groups are most easily separate along the first principal component, with 44% of the total variance being represented (**Figure 3d**). Although only 115 proteoforms could be quantified between the dissected tubules and glomeruli, these measurements were sufficient to cluster the samples into two clusters that aligned with their origin (**Figure 3e**). Notably, 20 of these proteoforms were differentially abundant with an FDR less than 0.01 **(Figure 3f)**.

**Figure 3:**
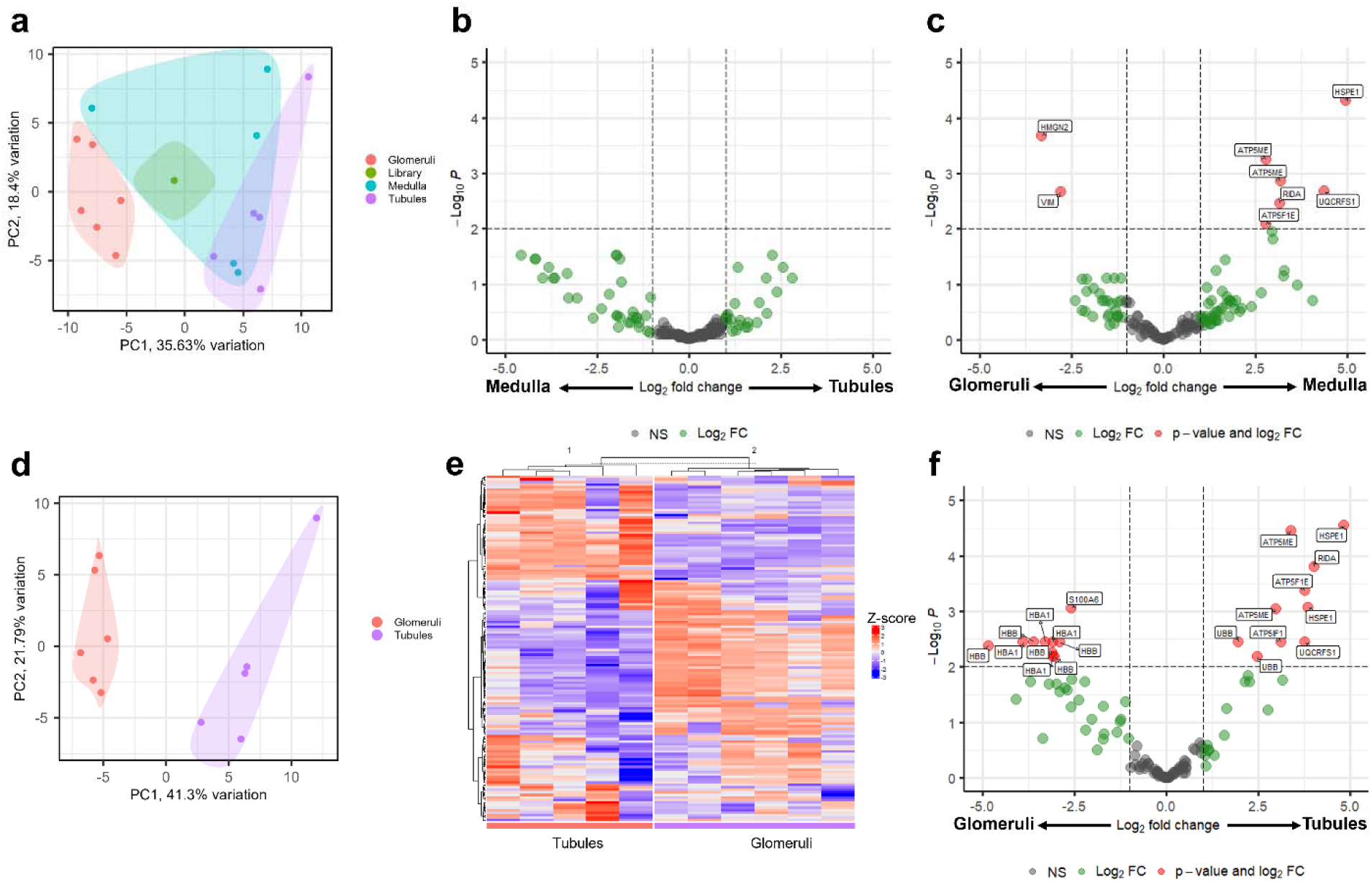
LCM-microPOTS-TDP analysis of human kidney FTUs. **(a)** Principal component analysis (PCA) of all replicates passing quality control derived from the three FTUs, as well as the aggregated library sample. **(b)** Volcano plot of proteoforms quantified between medulla regions (left) and tubules (right). **(c)** Volcano plot of proteoforms quantified between glomeruli (left) and medulla regions (right). **(d)** PCA of only tubule and glomeruli samples. **(e)** Heatmap of 115 proteoforms (y-axis) quantified between tubule and kidney samples (x-axis). Hierarchical clustering was applied to both proteoforms and samples. **(f)** Volcano plot of proteoforms quantified between tubules and glomeruli. Dotted lines for all volcano plots indicate thresholds for adjusted p-values < 0.01 and log_2_FC > 1 or < −1.

Of the proteoforms enriched in tubules, the majority (8 out of 10) were derived from mitochondrial proteins. This includes HSPE1, ATP5F1E, UQCRFS1, ATP5ME, and ATP5IF1. Many of these proteoforms are the full-length protein product, and in a few cases different proteoforms from the same protein showed similar trends in differential abundance between the two FTUs, such as with full-length HSPE1 and a M(Ox) HSPE1 proteoform, which both had log_2_ fold-changes greater than 3.8. On the other hand, proteoforms enriched in glomeruli were almost entirely derived from hemoglobin (9 out of 10). Interestingly, proximal and distal tubule rely on active transport mechanisms to reabsorb ions during filtration. Because tubules have higher energy demands, they contain more mitochondria than any other structure in the kidney, which aligns with our observations.^46^ Glomeruli contain bundles of capillaries and therefore would be expected to have more hemoglobin due to the increased concentration of red blood cells, again supporting our observations.

Taken together, these proteoform-level differences highlight known functional characteristics of these FTUs and provide validation of our quantitative approach with TopPIC processing. To aid in the production of these databases the library and dissected regions were searched with tandem use of the the TopPIC and TDPortal suite.^10^ This enabled roughly 33% increase in the number of annotations as demonstrated in a comparison of library overlap from an exemplary bulk top-down proteomics dataset presented in the Venn diagram in **Figure 4**. Indeed, the performance of a variety of TDP tools can be variable for the identification of proteoforms,^47^ and our approach for the automatic annotation and integration of MALDI-MSI relies heavily upon congruency between LCM-microPOTS-TDP and MALDI-MSI.

**Figure 4:**
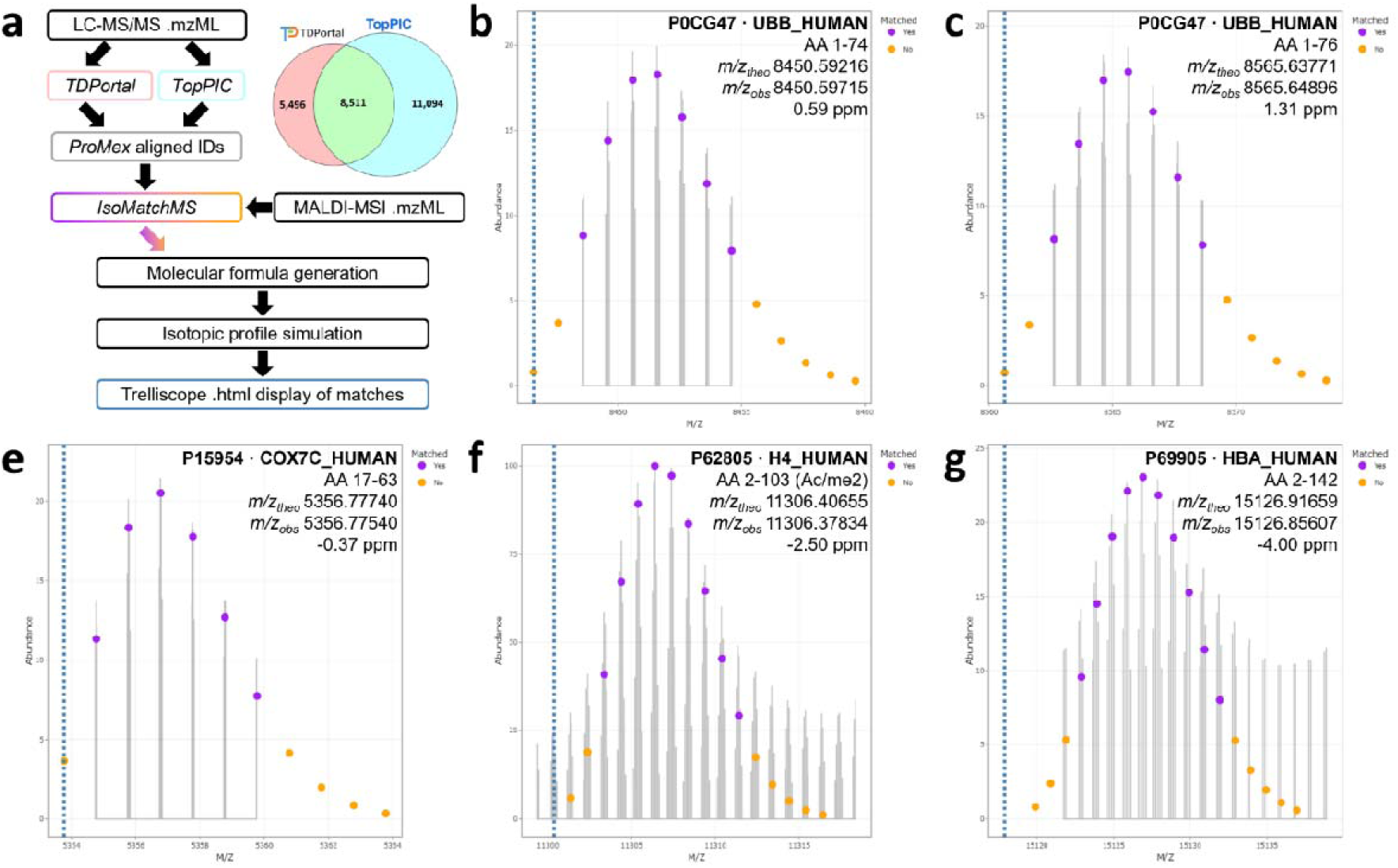
Bioinformatics workflow for annotation of MALDI-MSI datasets with LCM-microPOTS-TDP annotations. **(a)** Simplified schema of LC-MS/MS annotations pipeline where combined annotation using TDPortal and TopPIC enabled roughly 33 % more annotations as exemplified within the venn diagram from a bulk TDP dataset, ProForma strings from these aligned ProMex proteoform IDs are ingested into IsoMatchMS for isotopic profile matching and display within Trelliscope. Isotopic matching of **(b)** truncated ubiquitin B (1-74), **(c)** intact ubiquitin B (1-76), **(d)** truncated cytochrome c oxidase subunit 7C (17-63), **(e)** histone H4 (2-103) with a mass shift corresponding to an acetylation and dimethylation (+70.0418 Da), and **(f)** hemoglobin subunit A (2-142) are overlaid onto the averaged MALDI-MSI spectrum with annotation of matched and unmatched isotopes. For each respective window the UniProt accession ID, theoretical *m/z*, observed *m/z*, and calculated error is presented.

### Spatial Proteomics by MALDI-MSI within Human Kidney

Validation of any annotation for MALDI-MSI was often a tedious event and required manual matching and validation of isotopic profiles from proteins within the UniProt database. While many tools can seamlessly annotate TDP datasets,^47^ including those demonstrated above for processing of LCM-microPOTS-TDP data, these tools cannot accommodate simplistic MS1 level spectra generated by MALDI-MSI. To accomplish this, we utilized *IsoMatchMS*, a purpose-built program for MALDI-MSI annotation.^44^ As shown within **Figure 4** isotopic envelopes are automatically generated and visualized within trelliscope displays, which can then be sorted via Pearson correlation of the theoretical and observed isotopic profiles. While it is still highly recommended to visually inspect raw data, whether it be LC-MS/MS or MALDI-MSI, this level of automation facilitates higher throughput within our methodology critical for broad adoption and larger cohort studies.

Overall, from our LCM-microPOTS-TDP proteoform library containing 616 proteoforms, several dozen MALDI-MSI features could be annotated at the protein level. This is exemplified within **Figure 5**, where proteoforms are annotated by high resolution accurate mass with less than 3 ppm error,^48^ with subsequent confirmation of experimental isotopic profiles shown from the average MALDI-MSI spectrum within **Figure 4**. In the absence of an experimental proteoform database, ion images produced by MALDI-MSI are often matched to *in-silico* predicted intact masses of full-length proteins, whereere inclusion or exclusion of PTM or terminal truncations can easily lead to false identifications. Here, we demonstrate how our LCM-microPOTS-TDP database enabled facile identification of isomeric proteoforms, uncommon proteoforms, or truncated proteoforms. Notable annotations within MALDI-MSI include a truncated cytochrome c oxidase subunit 7C (17-63) **(Figure 5a)** and ubiquitin B (1-74) **(Figure 5b)**, and heavily modified positional isomers of histone H4 **(Figure 5f-h)**. This combination of a cellular-resolution imaging with a broad field of view enables the facile profiling of FTUs such as glomeruli within human tissues.

**Figure 5:**
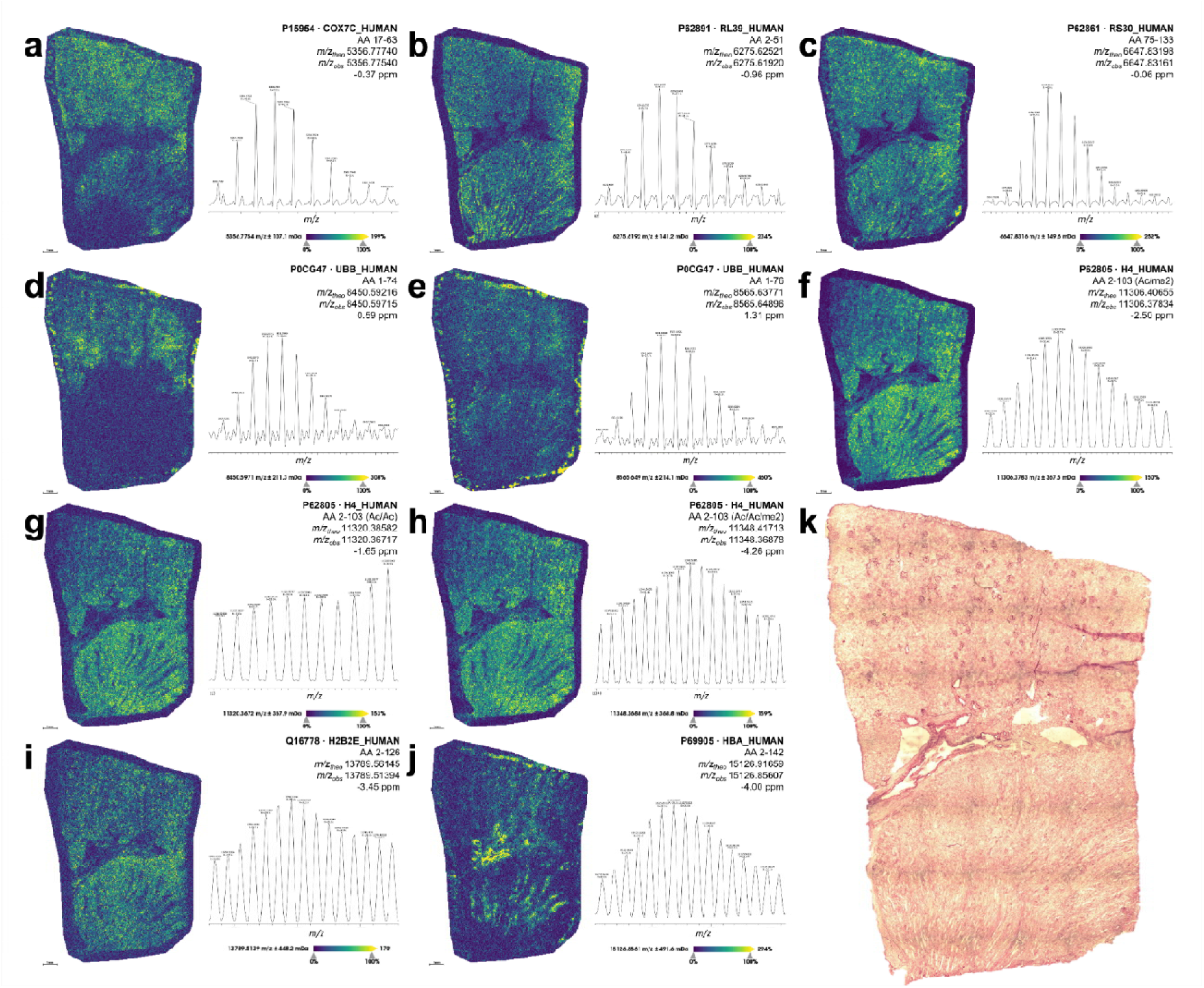
Ion images from baseline resolved isotopes of intact proteoforms with IsoMatchMS annotations. The UniProt accession ID, amino acid sequence, PTMs, theoretical and observed *m/z*, and mass measurement error are reported for **(a)** cytochrome c oxidase subunit 7C, mitochondrial, **(b)** large ribosomal subunit protein eL39, **(c)** small ribosomal subunit protein eS30, **(d)** truncated ubiquitin B, **(e)** intact ubiquitin B, **(f)** histone H4 acetylated and dimethylated (N-Ac/K20me2), **(g)** histone H4 diacetylated, **(h)** histone H4 diacetylated and dimethylated, **(i)** histone H2A type E, and **(j)** hemoglobin subunit A. The scale bar for **(a)** through **(j)** is 1 mm. **(k)** Shows a high-resolution image of the histological image taken post MALDI-MSI acquisition. Several hotspots can be noted off the tissue and were identified as embedding artifacts which did not influence analyses. **SMART** annotation:^43^ S (step size, spot size, total pixels) = 40 µm, 12 µm x 15 µm, 86,189 pixels; **M** (molecular confidence) = MS1, <5 ppm; **A** (annotations) = MS1 matching from LCM-microPOTS-TDP; **R** (resolving power) = 35k at *m/z* 11306; **T** (time of acquisition) = 736 min.

This high spatial resolution broad field-of-view analysis is well in line as an ideal case study, where preclinical and diseased samples can possess distinct spatial heterogeneity as previously demonstrated on sections containing tubular atrophy or a renal cell carcinoma.^13^ MALDI-MSI profiled several dozen glomeruli in this singular kidney tissue section. Although distinct proteoform markers were not found in the detected mass range localized solely to glomeruli, it was previously demonstrated larger proteoforms may be more region specific in kidney than the histone and other small proteoforms detected here.^14, 49^ Conversely smaller peptides, glycans, and metabolites also uniquely localize to these FTUs more readily.^19^

This absence of succinct proteoform markers from FTUs in our detected mass range for MALDI-MSI was not surprising given the inherent homogeneity of the proximal FTUs barring any acute or chronic conditions and the detection of primarily epigenetic markers in these analyses. Similarly, nearly ten-fold more proteoforms were annotated by LCM-microPOTS-TDP with 28 proteoforms being discriminate within pairwise comparisons between FTUs. Interestingly, we identified several ions with unique localizations between distinct regions (*e.g.*, cortex and medulla). For example, interfacial and medullary regions were densely abundant with hemoglobin subunits as depicted by MALDI-MSI, with disperse abundance within cortical regions **(Figure 5j)**. Within LCM-microPOTS-TDP, differential abundances of hemoglobin proteoforms (HBA/HBB) were found to be not significant between medullary and glomerular dissections **(Figure 3c)**, while glomeruli and tubular dissections did show significance **(Figure 3f)**.

Taken together, LCM-microPOTS-TDP and MALDI-MSI further contextualize each other, where the trends of high specificity and sensitivity LC-MS/MS are recapitulated within broad field-of-view imaging with several additional distinct localizations (which were not dissected **(Figure 5j)**. Furthermore, we identified a differentially abundant truncation of ubiquitin as highlighted within **Figure 6**. This annotation derived from the LCM-microPOTS-TDP database is the truncated form of ubiquitin (1-74) missing the C-terminal diglycine residues **(Figure 6a),** which is detected alongside the intact form (1-76) of ubiquitin **(Figure 6b)** and ubiquitin (UBB) was found to be differentially abundant within cortical FTUs, with significantly less abundance within glomeruli **(Figure 3f)**. This aligns with histological overlay and trends within ion images **(Figure 3d,e)**.

**Figure 6:**
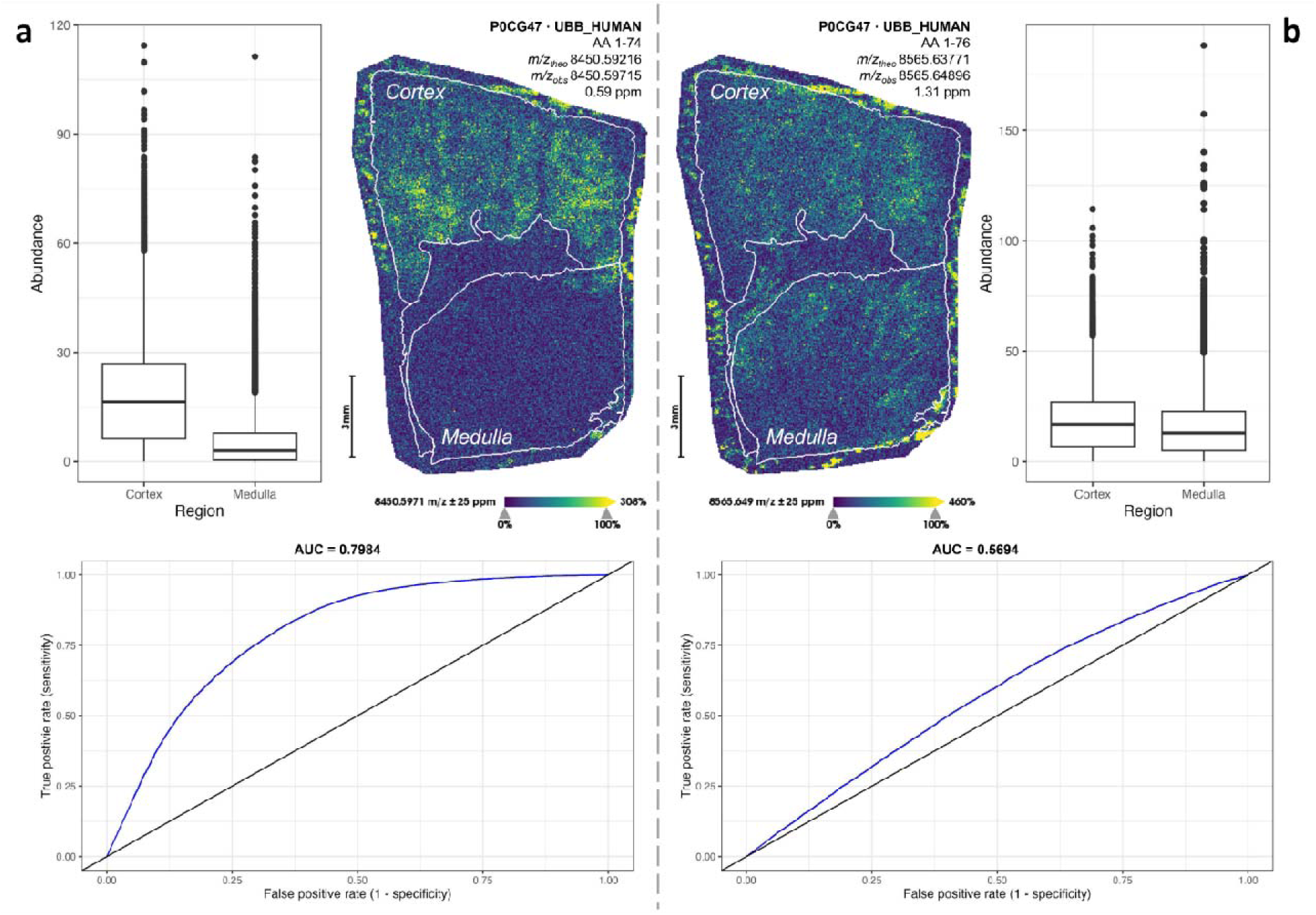
Detection of canonical and truncated forms of ubiquitin. The UniProt accession ID, amino aci sequence, PTMs, theoretical and observed *m/z*, and mass measurement error are reported for: **(a)** truncated ubiquiti B (1-74), which is presented with a box-and-whisker plot of normalized abundance within both the cortex and medulla, where the AUC within ROC analyses is 0.7984 showing increased abundance within the cortex; and **(b)** intact ubiquitin B (1-76), which is presented with a box-and-whisker plot of normalized abundance within both the cortex and medulla, where the AUC within discriminative analyses is 0.5694 showing no increased abundance within the cortex or medulla. Scale bars for **(a)** and **(b)** are 3 mm. Several hotspots can be noted off the tissue and were identified as embedding artifacts which did not influence analyses. **SMART** annotation:^43^ **S** (step size, spot size, total pixels) = 40 µm, 12 µm x 15 µm, 86,189 pixels; **M** (molecular confidence) = MS1, <5 ppm; **A** (annotations) = MS1 matching from LCM-microPOTS-TDP; **R** (resolving power) = 35k at *m/z* 11306; **T** (time of acquisition) = 736 min.

Interestingly, this truncated form of ubiquitin (1-74) appeared to have distinct localization in the cortex of the kidney. Indeed, receiver operating characteristic (ROC) analysis of this proteoform across these two areas provided an area under the curve (AUC) of 0.7984. Furthermore, ubiquitin (1-74) has been previously noted as a CXC chemokine receptor 4 (CXCR4) agonist,^50^ where CXCR4 has broad roles within cell proliferation, adherence, migration, and homing.^51^ While the significance and function of this unique localization of ubiquitin (1-74) within the cortex is unknown, production of ubiquitin (1-74) can be mediated through a rapid diglycine cleavage by insulin-degrading enzyme (IDE).^52^ At the mRNA level IDE is significantly enriched within the kidney and liver, and degradation of insulin is a well-established function of FTUs in the renal cortex.^53^ The integrated approach using both LCM-microPOTS-TDP and MALDI-MSI technology readily captured such protein processing events highlighting the potential for discovering proteoform biomarkers.

## Discussion

Here, we present an infrastructure for spatial TDP via the integration of LCM-microPOTS-TDP and MALDI-MSI. Although we (and others) have demonstrated significant gains in proteoform identifications in small amounts of tissue,^10, 27^ limitations still exist. For example, proteome coverage remains low relative to BUP based nanoPOTS or microPOTS format which routinely achieves thousands of protein identifications from a 10,000 µm^2^ dissected area.^54^ As we observed with our data, proteoform quantification precision and data completeness represent areas that can still be improved upon in future studies. Furthermore, the detection of proteoforms by MALDI-MSI has been practically limited to several dozen unique proteoforms with a high molecular mass limit of roughly 24 kDa.^12^ Other MSI capable techniques including nanospray desorption electrospray (nanoDESI) have recently expanded the upper molecular mass ceiling of proteoforms by MSI on healthy human kidney to beyond 70 kDa using highly sensitive charge detection MS.^14^ However, there is a consideration that many studies using nanoDESI-MSI detect highly abundant surface and cytoplasmic proteoforms,^15, 55^ where MALDI-MSI excels in detecting smaller regulatory and nuclear proteins, such as histones, at cellular resolution with throughput that enables several entire sections to be routinely imaged weekly. This leads to high complementarity, with specific use cases exemplified for each technological platform and vast potential for integration. As the technologies are rapidly evolving, and new instrumentation for intact protein analyses becomes available, we envision that each step within our workflow (e.g., sample preparation, data acquisition, and analyses) will be further improved to enable deeper proteoform coverage than what is currently attainable.

Our integrated LCM-microPOTS-TDP and MALDI-MSI approach on kidney also expands on previous efforts published from the Kidney Precision Medicine Project (KPMP), where the understanding of disease etiology and pathophysiology within human kidneys is a paramount goal.^19^ Previously, LCM-microPOTS-BUP enabled the identification of 92 unique proteins which were specific to either dissected proximal tubules or glomeruli with a total of 2,500 detected proteins from roughly 10 to 40 cells, which provided a roadmap for identifying the roles of these proteins in sub-structures of the renal corpuscle and nephron and validated affinity reagents in a non-targeted manner.^56^ In comparison, our TDP approach revealed 20 unique proteoforms, of which several peptides from each protein were previously detected from BUP, but were not found to be unique to these FTUs. All cell-type and FTU-type specific markers need additional validation, where biological variability (*i.e.*, age, ethnicity, gender, etc.) are additional factors which need to be controlled. Regardless, the detection of intact proteins demonstrates the potential of discovering new biology from “old” proteins.^57^ For example, even well characterized proteins, such as ubiquitin, have been identified in truncated forms with unique localization and unknown function within our imaging analyses

Regardless of caveats to various techniques, these approaches have found practical use within laboratories, and large consortia spanning from HuBMAP, where the focus has been on multiple organ systems,^34^ as well as more directed efforts focused upon specific organs such as the Lung Molecular Atlas Program (LungMAP)^58^, and the Cellular Senescence Network (SenNet)^59^ focused upon crucial transitions of cells throughout life. Altogether, efforts such as these are helping to build the Human Reference Atlas (HRA)^60^ and Human Protein Atlas (HPA)^61^, where proteoform informed analyses will be fundamental to deciphering complex biological mechanisms down to single cellular resolution and for advanced precision medicine;^3^ exemplifying that a fusion of proteomic practices is indeed the path forward for future functional proteomic studies. And further on the horizon, integration of these advanced proteomic technologies with other −omics assays will enable us to start deciphering fundamental principles of biology.

When considering broad applications of these techniques, there are clearly upfront investments which are paramount to success. A persistent burden of method optimization for all spatial proteomics demands a rigorous optimization of sample washes for each new tissue type analyzed limited only by the creativity of the user,^62^ and assay transferability and detection of targeted proteoforms between organs or organisms cannot be guaranteed. Additionally, advanced LCM approaches are now breaking into the subcellular regime,^63^ and traditional methods employing high-end MS platforms for TDP do not provide the sensitivity to bridge to this resolution. The next decade will provide exciting developments such as the translation of charge detection to chromatographic time scales,^64^ and new instrumentation to further characterize proteoforms.^65^ Yet, our current approach can still be broadly applied to profile many signal peptides, hormones, and other small proteoforms with high biological significance in many human tissues.

## Conclusions

The data generated through the integration of LCM-microPOTS-TDP and MALDI-MSI enabled the bridging of several hundred high confidence proteoform annotations to several dozen direct near-cellular resolution proteoform maps. LCM-microPOTS-TDP provided a narrow field-of-view focusing on several FTUs and identifying 20 discriminate proteoforms between glomeruli and tubules relevant to the known physiological processes within the kidney. This analysis enabled the production of a library of 616 proteoforms, showing high diversity of proteoforms within the renal corpuscle and nephron as the FTUs cross distinct anatomical regions. Tandem application of MALDI-MSI demonstrated the ability to broadly profile an entire tissue section within a practical period at near cellular resolution, focusing upon the relative quantitation of distinct cortical and medullary markers. Together these datasets offer unique insights into the physiological processes of the kidney, such as the distinct localization of proteoform truncation events.

## Declarations

### Ethics approval and consent to participate

We have complied with all ethical regulations related to this study. All experiments on human samples followed all relevant guidelines and regulations at host institutions and informed consent for remnant tissue collection was acquired in accordance to Institutional Review Board policies at BIOMIC at Vanderbilt University, a Tissue Mapping Center associated with the HuBMAP.

### Consent for publication

Not applicable.

### Availability of data and materials

The datasets supporting the conclusions of this article are available both on the HuBMAP data portal, [https://doi.org/10.35079/HBM334.DQWS.354 https://doi.org/10.35079/HBM666.BBVZ.767], as well as on MassIVE [MSV000093722, https://massive.ucsd.edu/ProteoSAFe/dataset.jsp?task=f7afef33eea446d99f72f3220ef53a64].

### Authors’ contributions

KJZ, JMF, and RK wrote the original draft, where KJZ and JMF finalized formal analyses and discussion. KJZ completed all mass spectrometry imaging analyses, where KJZ and RK analyzed resulting datasets. DJD and LAL processed top-down annotations for mass spectrometry imaging, and DJD performed statistical analyses on these datasets. KJZ completed all imaging visualization. MV and DV processed all laser capture microdissection and microscopy analyses, where YCL, MZ, YZ, JMF, and LPT developed top-down proteomics methodology. JMF, MZ, and SMW completed all liquid chromatography analyses, and JMF processed all top-down datasets and completed all top-down visualization. MZ administered the project, where LPT, YZ, and MZ conceptualized the project, and LPT acquired funding and provided supervision. All contributed to final editing of the manuscript and all authors read and approved the final manuscript.

## Acknowledgements

The authors would like to acknowledge Jamie Allen and Dr. Jeffrey Spraggins at Vanderbilt University for providing the kidney tissue sections through the HuBMAP Consortium. Dr. Mikhail Belov at Spectroglyph, LLC, as well as Dr. Gordon Anderson and Chris Anderson at GAA Custom Electronics, LLC, for technical support for the MALDI source. The authors would also like to acknowledge Drs. Tobias Wörner, Kyle Fort, Maria Reinhardt-Szyba, and Alexander Makarov of Thermo Fisher Scientific for technical guidance and licensing of the instrument. Finally, we thank Dr. Neil Kelleher, Ryan Fellers, and Joseph Greer from Northwestern University for access and assistance with TDPortal and Matthew Monroe from Pacific Northwest National Laboratory for assist with MassIVE upload.

