## Supplementary Information for "Spatial top-down proteomics for the functional characterization of human kidney"

**Table of Contents:**

**Figure S1:** Box plots showing coefficients of variation for each functional tissue unit

**Figure S2:** Bar plots showing data completeness for each functional tissue unit

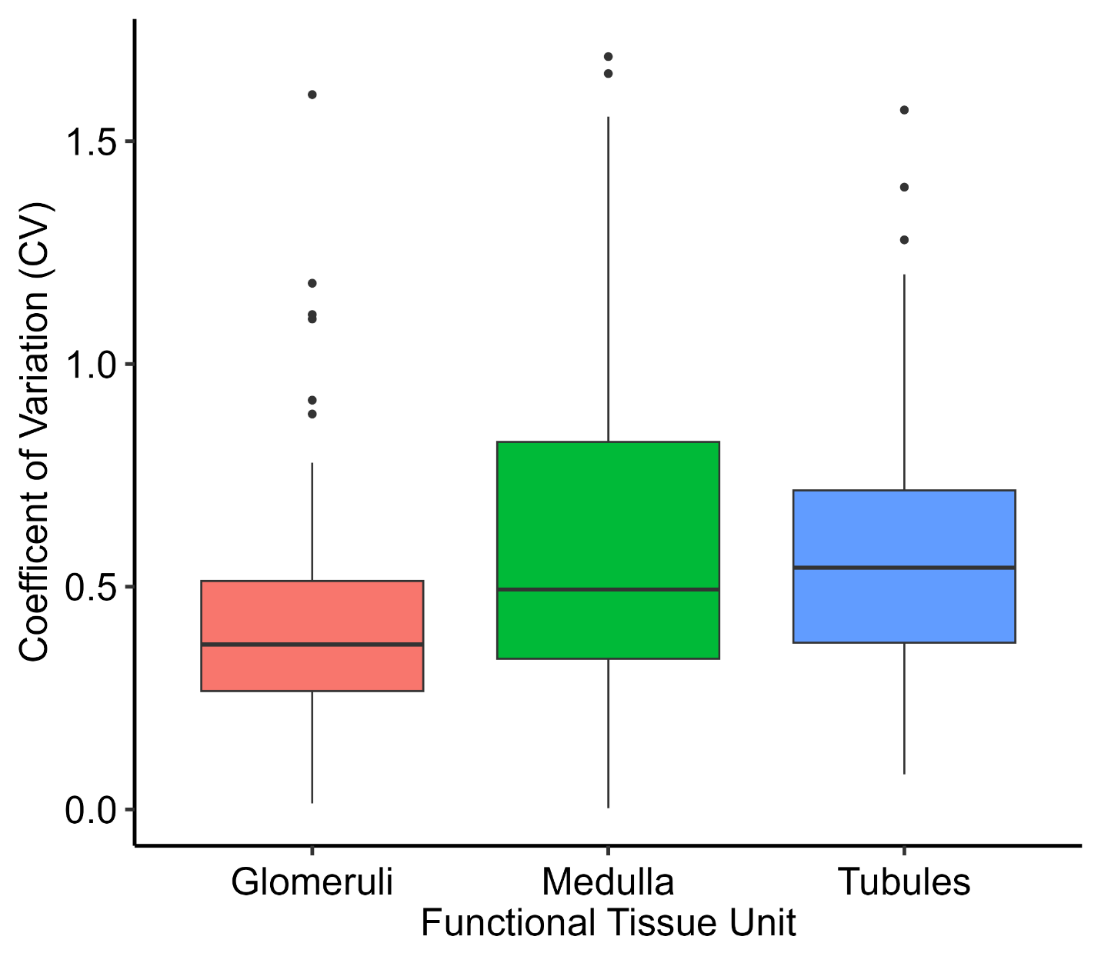

**Figure S1:** Distributions of coefficients of variation (CV) for each functional tissue unit.

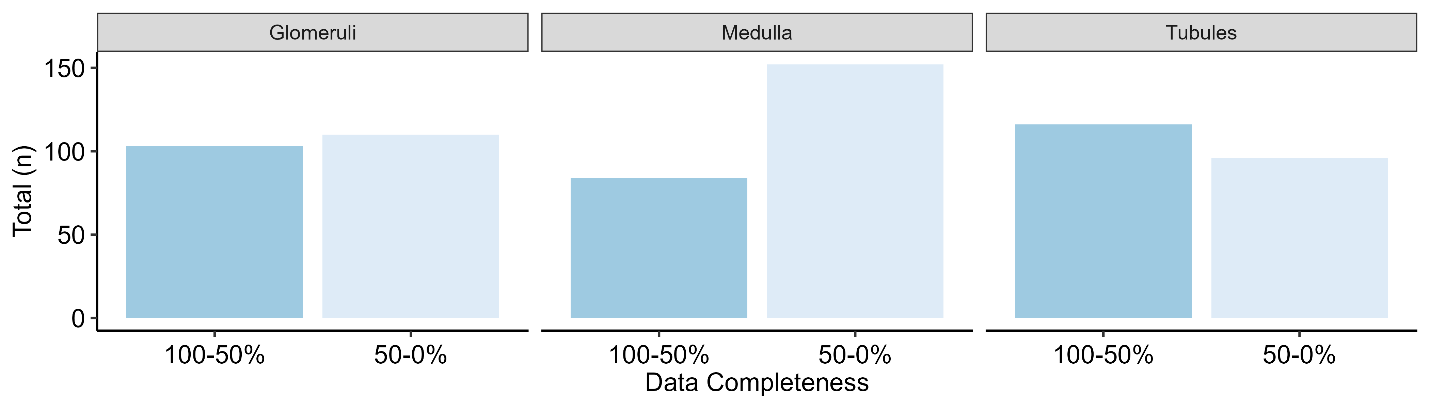

**Figure S2:** Bar plots showing data completeness for each functional tissue unit. Proteoforms were separated into two bins depending on if data completeness was greater than or less than 50%.
